# Occupation and Parkinson disease in Women’s Health Initiative Observational Study

**DOI:** 10.1101/535732

**Authors:** Igor Burstyn, Andrea Z. LaCroix, Irene Litvan, Robert B. Wallace, Harvey Checkoway

## Abstract

**Introduction:** There is a lack of consistency in associations between workplace factors and risk of Parkinson disease (PD), and paucity of such data on women. We took a classic occupational epidemiology approach that assesses associations with occupational groups in order to derive insights about potential occupation-specific exposures that may be causal.

**Methods:** The Women’s Health Initiative (WHI-OS) is a prospective cohort that enrolled 91,627 postmenopausal women, 50 to 79 years of age from 10/01/93 to 12/31/98, at 40 clinical centers across the US with average follow-up of 11 years, who reported up to three paid jobs held the longest since age 18; these jobs were coded and duration of employment calculated. We defined the case by self-report of doctor-diagnosed PD (at baseline or follow-up), death attributed to PD, or medication consistent with PD.

**Results:** Among 2,590 cases, we report evidence of excess risk among “counselors, social workers, and other community and social service specialists”. There was a suggestion of increase in risk among post-secondary teachers, and “building and grounds cleaning and maintenance”. There was also evidence of deficit in risk among women who worked in sales.

Results with ever-employed and duration were similar, except for evidence of excess of risk among “health technologists and technicians” with more than 20 years of employment. Longer duration of life on a farm was associated with higher risk.

**Conclusion:** Our findings paint a largely reassuring picture of occupational risks for PD among US women, especially for trades largely unaffected by recent technological advances.

## Introduction

There is epidemiologic evidence that some occupational exposures are etiologically related to Parkinson disease (PD) [1 2]. There is consistent supportive evidence for manganese and some pesticides. Among pesticides, the strongest etiologic evidence is for paraquat and maneb used in agricultural settings [2 3]. For other occupational exposures, such as shiftwork (light-at-night) and solvents, the evidence is mixed, despite some biological plausibility [1]. There are suggestive findings for trichloroethylene, but no other specific solvents [4]. Excesses of PD was observed in various occupations with mixed findings [5-14], few of which are specific to women except for excess among “assistant nurses” [7], teachers [12], and a report that “complex work with people” is associated with PD only in women [15]. Occupations associated with a higher socio-economic status in the US tend to show excess risk of PD mortality [16]. Most prior research on occupational risk for PD has been conducted among men, who historically have been the predominant workforce in occupations known or presumed to be hazardous. However, such evidence cannot be assumed to equally apply to women, because job types have differed between men and women historically. Furthermore, there is a concern that even in the same job, actual exposures may differ between men and women due to variation in tasks and materials [17 18]. More importantly, most research has focused on men, and given that women may be vulnerable to the same exposures, research on female workers is warranted to test coherence of evidence. Consequently, it is premature to limit research into etiology of PD among women on a specific workplace exposures. Instead, our goal is to take a classic occupational epidemiology approach that first assesses associations with occupational groups, in order to derive insights about potential occupation-specific exposures that may be causal. Our main objective is to estimate the risk of PD in association with common occupations in the (exclusively female) participants in the observational arm of the Women’s Health Initiative (WHI-OS) cohort.

## Methods and Materials

### Cohort

The WHI-OS is a prospective cohort that enrolled 93,676 postmenopausal women, 50 to 79 years of age from October 1, 1993, to December 31, 1998, at 40 clinical centers across the US [19-21]. Eligibility criteria included 50 to 79 years of age, postmenopausal status, willingness to provide informed consent, and at least a 3-year life expectancy. A total of 4.1% participants were lost to follow-up or had stopped follow-up during the study and nearly 94% participants provided responses each year over an average 11.4 years. Participants updated medical history annually through mailed self-administered or telephone-administered questionnaires. Annual mailed questionnaires elicited medical history, including diagnosis of PD and use of medications, with about 95% response rates. Death was ascertained through clinical center follow-up of family reports and checks with the National Death Index. We used version of the data available in October 2017. Human subjects review committees at all participating sites approved WHI study protocols. The proposed research has been reviewed and approved by the Publications and Presentations Committee of the WHI and reviewed by Drexel University IRB.

### Outcome Ascertainment

We defined a case of PD if a physician-diagnosed PD was reported at either baseline (form F30), or any subsequent years by self-report (forms 33, 134, 143, 144, 145, 147, 148), if death was attributed to PD (forms 120, 124 February 28, 2017), or if medication consistent with a PD diagnosis was reported on either form 44 (September 12, 2005) or 153 (September 30, 2015). Self-report of PD correlates well with physician diagnosis: in the Women’s Health and Aging Study I, self-report of PD had perfect specificity and 89% sensitivity compared to record review [22]. Using a PD questionnaire examining the utility of screening questions compared to medical record diagnosis in clinic patients and controls in Rochester, the questionnaire also had perfect specificity, and 89% sensitivity [23]. The pharmacologic signature for PD is highly specific [24] and we interpreted any report of use of either non-ergoline dopamine receptor agonists or levodopa with or without decarboxylase inhibitors as evidence of PD diagnosis. Cases of PD identified only by medication were reviewed to confirm coherence with PD by I.L.; WHI lacked data on dosage of medication that would allow us to exclude cases with certainty. Date of onset or diagnosis of PD in the WHI is unknown.

### Exposure Assessment

At baseline, 91,627 women (97.8%) responded to the occupational history questionnaire (form 42, Q26.1-26.3). Women reported up to three paid jobs (full-time or part-time) held the longest since age 18. For each job, women reported the job title and industry, as well as the age at which work began and total duration of employment. We selected members of the WHI-OS cohort who had at least one paid job outside of the home to control for biases that arise from selection into the workforce. In approaching questionable entries in occupational histories, we elected to exclude them rather than impute, owning to large sample size available to us and to limit bias from assumptions inherent in imputations. We excluded women when (a) at least one of the jobs appears to have lasted >100 years; (b) at least one job reported to have started before age of 18; (c) at least one job reported to have started after interview; (d) at least one job reported to have ended at age that is 10 years older than age at interview.

Job histories were coded by the best available (trained) coders into standard categories of occupation used by the US Census by staff of the US National Institute for Occupational Safety and Health (NIOSH) (i.e. gold standard coding and independent of knowledge of any of the health outcomes in WHI). Of the 274,881 records of occupational history, 48,908 were removed for coding purposes because both job title and industry were blank. The rest (225,973 records) were divided into 46 files (5,000 observations each in 45 files, and 973 records in the last file). Some of these records contained no useful information and were recorded as missing. Files were first processed in the NIOSH Industry and Occupation Computerized Coding System (NIOCCS) for auto-coding, next sent to trained coders for computer-assisted coding, then to the quality control (QC) coder for review [25]. NIOCCS confidence level was set to >90% confidence of accuracy, leading to auto-coding of 45.3% of occupations. Six previously trained coders received a refresher course that introduced the new 2010 census coding scheme and reviewed NIOCCS. Each coder received a post-auto-coded file of 5,000 records. If codes had not been assigned, the coder used the NIOCCS computer-assisted program to do so. The first three files were reviewed by QC coder 100%, then a 50% random sample were reviewed in files 4-14, and a 25% random sample in the rest. Because systematic errors were corrected in the first 3 files, the coders were retrained; since then, no systematic errors were detected. NIOCCS (both auto-coding and computer-assisted coder coding) first assigned the census 2010 occupation codes, then the corresponding Standard Occupational Classification (SOC) 2010 codes [26] were determined using the crosswalk built into NIOCCS. Because SOC 2010 is more detailed than the 2010 census, some records have multiple SOC values. Separate codes were created for non-paid workers and military occupations.

We created a variable for each occupation that represents having ever worked in that occupation; if multiple occupations were coded, then we allowed multiple classifications. Duration in each occupation was calculated based on the years a woman started and stopped working at each job, summed for each woman across all jobs with the same SOC 2010. During creation of ever employed variables, we eliminated jobs with no SOC 2010 codes and no information on duration of work. Because some SOC 2010 contained few PD cases, we aggregated SOC hierarchically till the number of cases ever-employed in an occupation was at least 100, except that we did not aggregate beyond 2-dig SOC 2010 regardless of number of exposed cases to preserve interpretability. Duration of employment in a given occupation was only examined for occupations with at least 100 cases “exposed to an occupation”.

### Statistical Analysis

We compared the distributions of demographic and lifestyle covariates among PD cases versus non-cases to assess the potential for confounding. We estimated relative risks (RR) and associated 95% confidence intervals (CI) via Poisson regression with robust variance estimate [27]. We examined each occupation (ever-employed, years employed in pre-specified categories) separately with and without adjustment for potential confounders (cubic polynomial of age at baseline, years of living on a farm, pack-years of smoking, coffee and alcohol consumption, region of the US, ethnicity, education, income, marital status, reported never employed) selected *a priori* [28]. Ever-employed variables had no missing values, but could have deficient information on duration; we retained such records as “unknown” in analysis by duration. We classified missing value of a covariates into a separate category in regression analysis, instead of excluding associated records. All statistical analysis and data management were implemented in SAS 9.4 (SAS Institute, Cary, NC).

## Results

After excluding women with errors in description of occupational history, we retained 80,646 subjects (88% of those who responded to occupational questionnaire), along with 2,590 cases of PD, a 3.21% rate (Table I). Only 186 cases were recorded at baseline, contemporaneously with occupational histories. The selected cohort exhibits patterns of increase chance of the diagnosis with age, regional and ethnic variation, as well as excess risk with longer farm living (which remained after adjustment for work-related factors, detailed below). Consumption of coffee tended to confer reduced risk. Pack-years of smoking did not appear to be associated with the risk in a monotonic manner. There was no pronounced variation by marital status, family income and education, lowering concerns about confounding by socio-economic factors.

**Table I:**
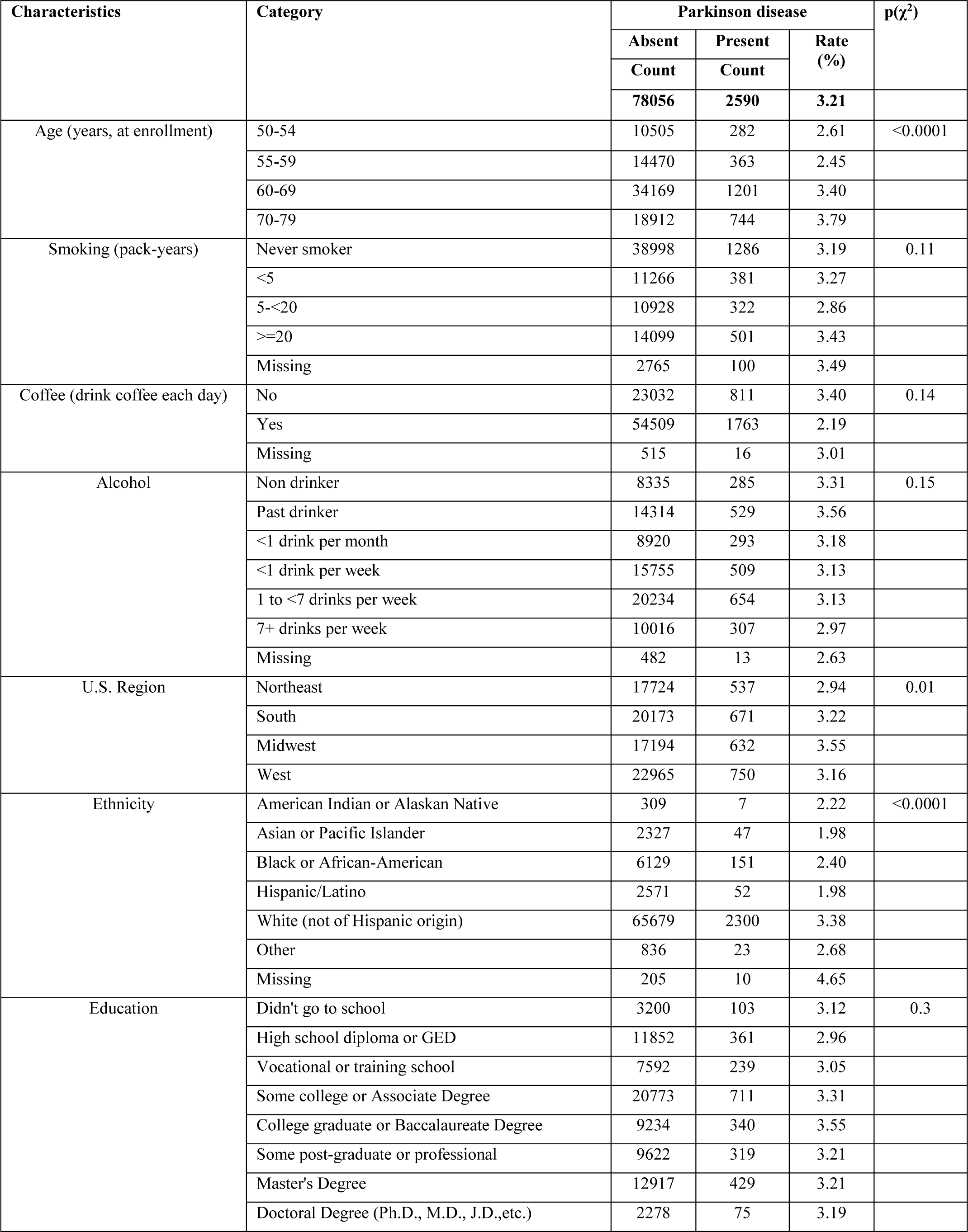
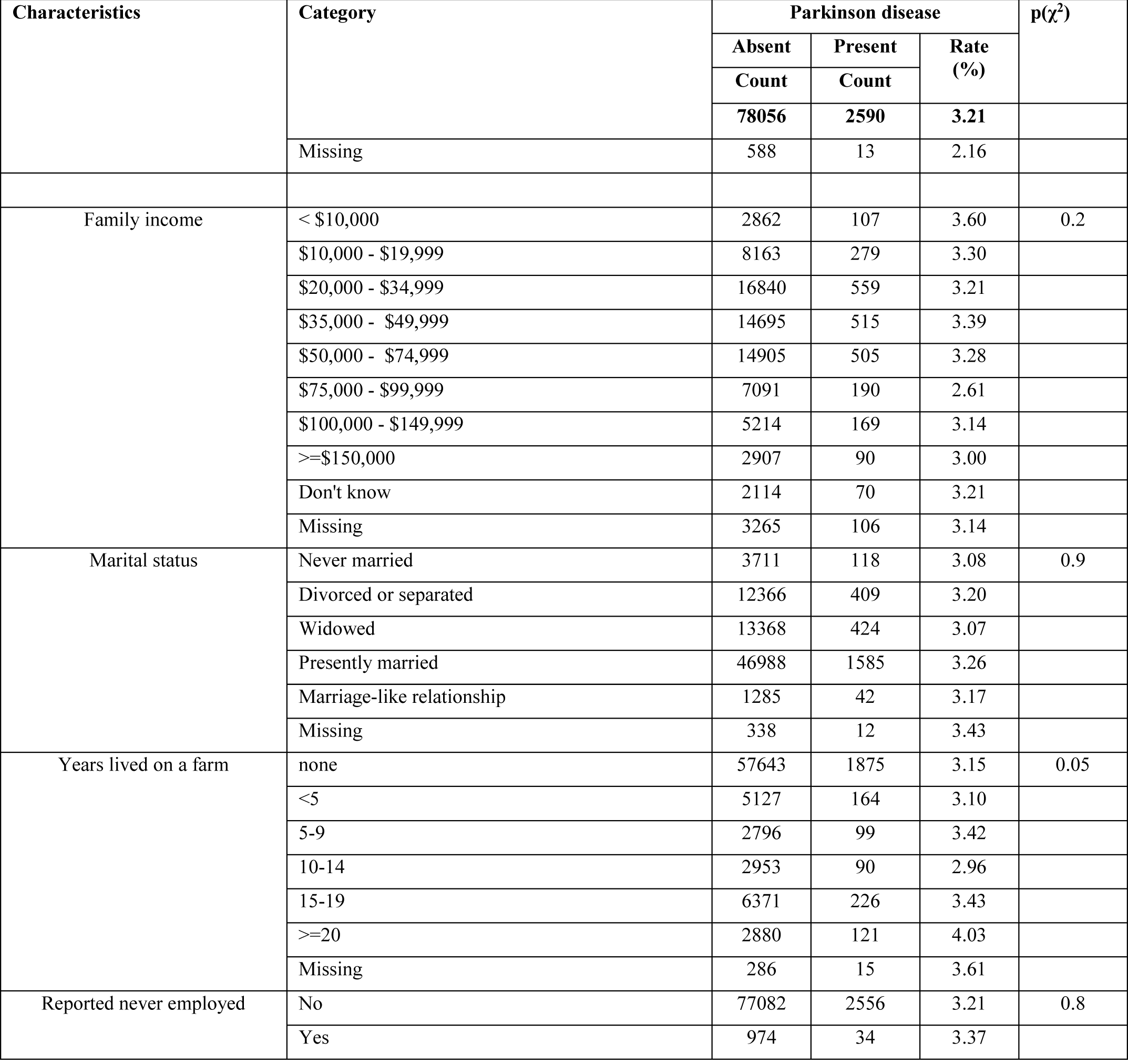
Cases and non-cases of Parkinson disease in portion of Women’s Health Initiative Observational Study with complete occupational histories

Distribution of cases, crude RR and adjusted RR by having ever been employed among 3 reported jobs in a specific occupation is displayed in Table II. There was evidence of excess risk of PD among women who worked as “counselors, social workers, and other community and social service specialists”, with 175 exposed cases and adjusted RR 1.18 (95% CI 1.01, 1.38). There was a suggestion of average increase in risk among top executives (48 exposed cases, adjusted RR 1.18 (95%CI 0.89, 1.56)), post-secondary teachers (104 exposed cases, adjusted RR 1.17 (95%CI 0.95, 1.45)), and “building and grounds cleaning and maintenance” (53 exposed cases, adjusted RR 1.21 (95%CI 0.92, 1.60)). There was also evidence of deficit in risk among women who worked in sales, especially retail. These effect estimates required adjustment for covariates to reveal evidence of an association. There was no evidence of association with farming, fishing and forestry, but there were only 7 exposed cases.

**Table II:** Having ever been employed in specific occupations and risk of Parkinson disease in portion of Women’s Health Initiative Observational Study with complete occupational histories (80,646 women)

| Occupation (SOC 2010 code) | Parkinson Disease |  |  | Estimated relative risk (95%CI) |  |  |  |
| --- | --- | --- | --- | --- | --- | --- | --- |
|  | Cases |  | Non-cases | Unadjusted |  | Adjusted for factors in Table I |  |
|  | N | % exposed | N |  |  |  |  |
| Top executives (11-1000) | 48 | 3.63 | 1276 | 1.00 | (0.99, 1.02) | 1.18 | (0.89, 1.56) |
| Advertising, marketing, promotions, public relations, and sales managers (11-2000) | 18 | 2.55 | 689 | 0.99 | (0.98, 1.01) | 0.82 | (0.52, 1.30) |
| Operations specialties managers (11-3000) | 44 | 2.95 | 1446 | 1.00 | (0.99, 1.01) | 1.06 | (0.70, 1.26) |
| Other management (11-9000) | 277 | 3.39 | 7895 | 1.00 | (1.00, 1.01) | 1.09 | (0.96, 1.23) |
| Business operations specialists (13-1000) | 114 | 3.23 | 3411 | 1.00 | (0.99, 1.01) | 1.03 | (0.85, 1.24) |
| Financial specialists (13-2000) | 86 | 3.02 | 2759 | 1.00 | (0.99, 1.00) | 0.93 | (0.75, 1.15) |
| Computer, math, architecture, engineering (15-1100, 15-2000, 17-000) | 75 | 3.10 | 2343 | 1.00 | (0.99, 1.01) | 0.98 | (0.78, 1.23) |
| Life, physical, and social science (19-0000, 19-4000) | 92 | 3.51 | 2531 | 1.00 | (1.00, 1.01) | 1.08 | (0.87, 1.34) |
| Counselors, social workers, and other community and social service specialists (21-1000) | 175 | 3.73 | 4513 | 1.00 | (1.00, 1.01) | 1.18 | (1.01, 1.38) |
| Legal (23-0000) | 31 | 3.03 | 993 | 1.00 | (0.99, 1.01) | 0.98 | (0.68, 1.40) |
| Preschool, primary, secondary, and special education school teachers (25-2000) | 522 | 3.26 | 15487 | 1.001 | (0.998, 1.004) | 0.98 | (0.88, 1.10) |
| Other teachers, librarians (25-3000, 25-4000, 25-9000) | 168 | 3.21 | 5070 | 1.00 | (0.99, 1.01) | 0.99 | (0.85, 1.16) |
| Postsecondary teachers (25-1000) | 104 | 3.76 | 2660 | 1.01 | (1.00, 1.01) | 1.17 | (0.95, 1.45) |
| Arts, design, entertainment, sports, and media (27-0000) | 134 | 3.11 | 4171 | 1.00 | (0.99, 1.00) | 0.93 | (0.79, 1.11) |
| Health diagnosing and treating practitioners (except RN) (29-1000) | 56 | 3.07 | 1768 | 1.00 | (0.99, 1.01) | 0.93 | (0.72, 1.21) |
| Registered nurses (29-1141) | 213 | 3.48 | 5915 | 1.00 | (1.00, 1.01) | 1.07 | (0.93, 1.23) |
| Health technologists and technicians, other healthcare (29-2000, 29-9000) | 105 | 3.47 | 2925 | 1.00 | (1.00, 1.01) | 1.10 | (0.90, 1.33) |
| Healthcare support (31-0000) | 104 | 3.16 | 3184 | 1.00 | (0.99, 1.01) | 1.03 | (0.85, 1.25) |
| Protective service (33-0000) | 22 | 3.44 | 618 | 1.00 | (0.99, 1.02) | 1.13 | (0.75, 1.71) |
| Food preparation and serving related (35-0000) | 129 | 3.08 | 4059 | 1.00 | (0.99, 1.00) | 0.97 | (0.81, 1.15) |
| Building and grounds cleaning and maintenance (37-0000) | 53 | 3.46 | 1481 | 1.00 | (0.99, 1.01) | 1.21 | (0.92, 1.60) |
| Personal care and service (39-0000) | 112 | 3.00 | 3623 | 1.00 | (0.99, 1.00) | 0.96 | (0.79, 1.15) |
| Sales (other) (41-0000 except 41-2031) | 279 | 2.89 | 9375 | 1.00 | (0.99, 1.00) | 0.89 | (0.79, 1.01) |
| Retail salespersons (41-2031) | 202 | 2.87 | 6843 | 1.00 | (0.99, 1.00) | 0.87 | (0.75, 1.00) |
| Financial clerks (43-3000) | 222 | 3.02 | 7138 | 1.00 | (0.99, 1.00) | 0.91 | (0.80, 1.05) |
| Information and record clerks (43-4000) | 199 | 2.93 | 6591 | 1.00 | (0.99, 1.00) | 0.91 | (0.79, 1.05) |
| Material recording, scheduling, dispatching, and distributing workers (43-5000) | 47 | 2.57 | 1784 | 0.99 | (0.99, 1.00) | 0.81 | (0.61, 1.07) |
| Secretaries and administrative assistants (43-6010) | 568 | 3.30 | 16661 | 1.001 | (0.998, 1.004) | 1.02 | (0.93, 1.13) |
| Other office and administrative support workers (43-9000) | 404 | 3.28 | 11905 | 1.001 | (0.997, 1.004) | 1.03 | (0.92, 1.14) |
| Supervisors of office and administrative support workers, communications equipment operators (43-1000, 43-2000) | 98 | 2.87 | 3322 | 1.00 | (0.99, 1.00) | 0.90 | (0.73, 1.10) |
| Farming, fishing, and forestry (45-0000) | 7 | 3.35 | 202 | 1.00 | (0.98, 1.03) | 1.04 | (0.50, 2.17) |
| Construction and extraction (47-0000) | 8 | 2.38 | 328 | 0.99 | (0.98, 1.01) | 0.74 | (0.37, 1.47) |
| Installation, maintenance, and repair (49-0000) | 11 | 3.13 | 341 | 1.00 | (0.98, 1.02) | 0.99 | (0.55, 1.77) |
| Production (51-0000) | 155 | 2.88 | 5227 | 1.00 | (0.99, 1.00) | 0.89 | (0.76, 1.06) |
| Transportation and material moving (53-0000) | 44 | 2.63 | 1632 | 0.99 | (0.99, 1.00) | 0.84 | (0.63, 1.13) |

Estimated associations with duration of work in selected occupations with sufficient number of cases are displayed in Table III; only adjusted RR are given as they were not materially different from crude ones (crude estimates are available upon request). Again, there is evidence of trend towards higher risk among “counselors, social workers, and other community and social service specialists”, with risk increasing from 3.47% to 3.91% with 1-5 years of work versus more than 20; relative to unexposed, the average estimated RR show monotonic trend of 1.09, 1.15, 1.21, 1.27 for 1-5, 5-10, 10-20 and >20 years, respectively. We note evidence of excess of risk among “health technologists and technicians” with more than 20 years of employment, based on 39 exposed cases, RR 1.46 (95%CI 1.07, 1.99). Women who were post-secondary teachers for more than 10 years tended to be at higher risk as well: RR 1.33 (95%CI 0.92, 1.92) and RR 1.23 (95%CI 0.84, 1.81) for 10-20 and 20+ years of work, respectively (29 exposed cases in each category). However, for these two occupation there was no consistent evidence of trend across all categories of duration. There is further evidence of reduced risk among retail salespersons, especially with more than 20 years of work: 13 exposed cases, RR 0.71 (95%CI 0.41, 1.22). Analyses that use duration as a continuous variable did not reveal patterns that add to information contained in figures presented above, and are therefore not shown.

**Table III:**
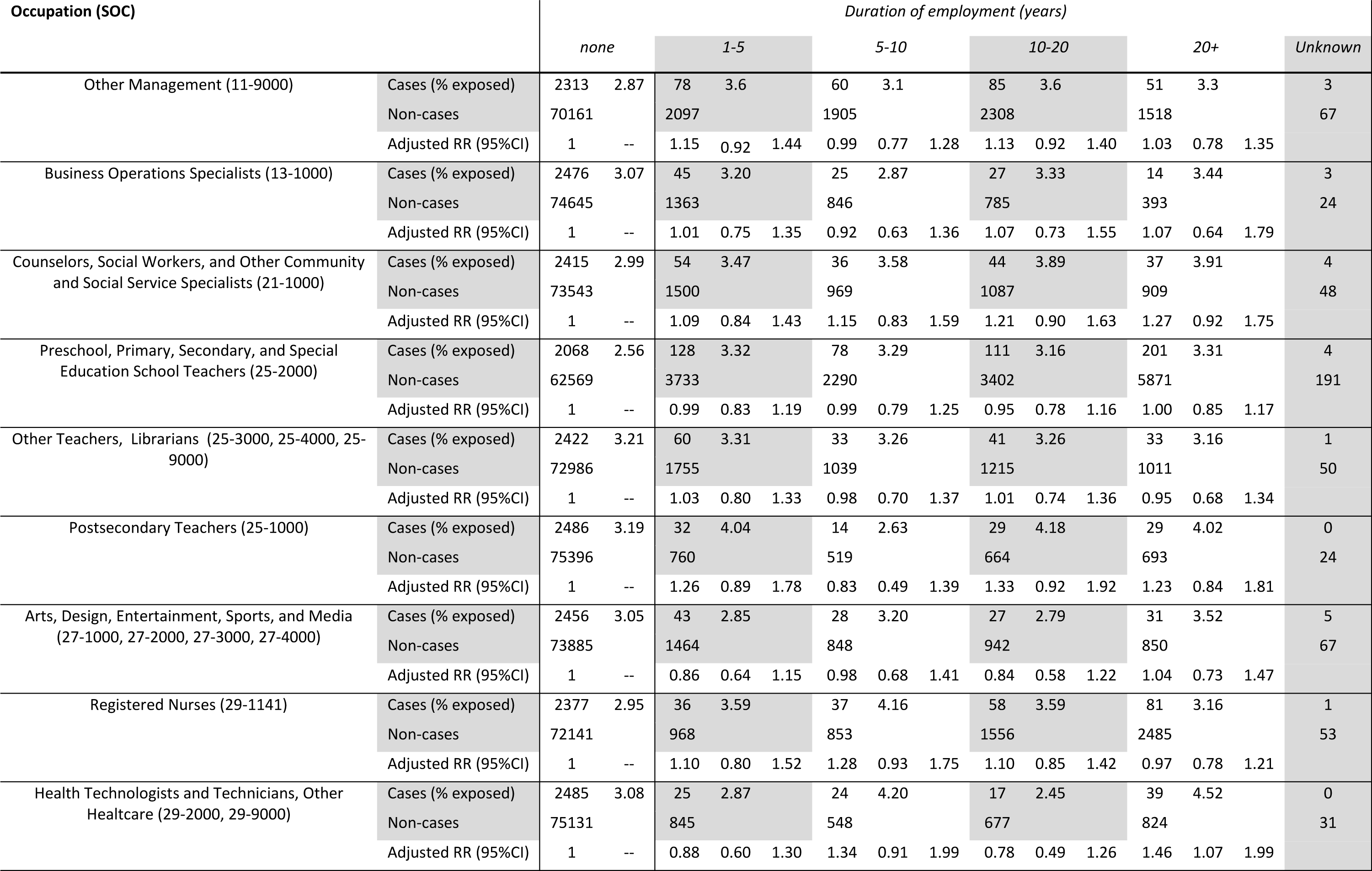
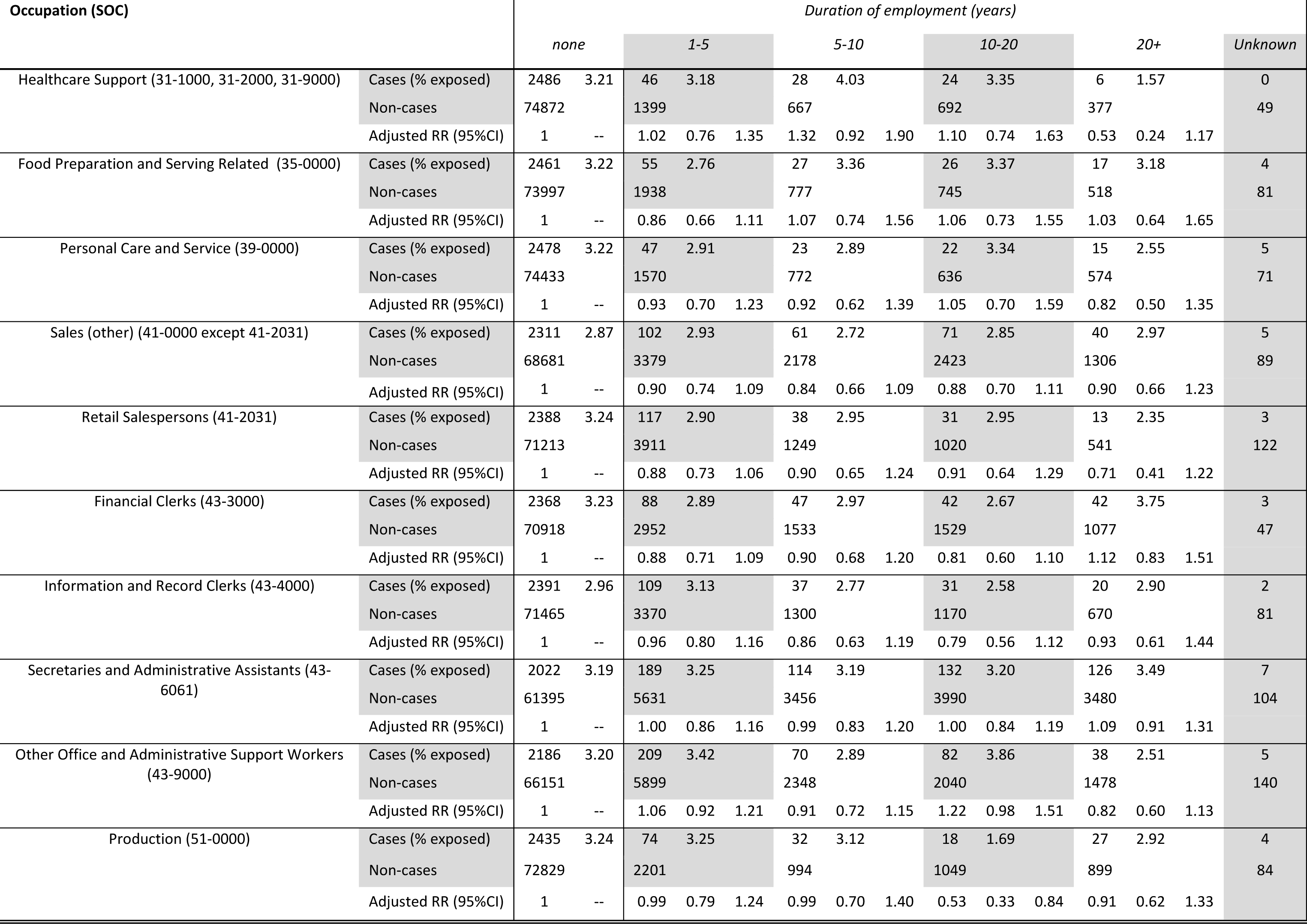
Duration of employment and risk of Parkinson disease in portion of Women’s Health Initiative Observational Study with complete occupational histories (80,646 women); adjusted models account for covariates in Table I.

## Discussion

We present result of analysis of diverse cohort of US women with the number of cases that exceeds even the largest proportional mortality study (2,293 mostly white women) [6]. The majority of PD cases (>90%) were ascertained during prospective follow-up after information on occupational history was recorded. Information on occupational history was detailed enough to identify occupation and duration of work for 88% of the cohort for a substantial portion of expected working life (by assessment of up to three longest-held jobs). We considered a comprehensive list of plausible confounders (including socioeconomic factors known to affect occupational associations with PD [16]), and, although residual confounding can never be ruled out, we minimized it. As such, our analysis is among the most informative regarding impact of occupation among US women and risk of PD.

Our results generally bolster evidence of historic investigation that linked work in healthcare, teaching and social work to the elevated risk of PD [5]; however, there are some apparent inconsistencies. The only “assistant nurses” were at elevated risk of PD among Swedish women in a record-linkage study of hospital discharge [7], but none of the nursing occupations in our work showed excess risk, unlike health technologists and technicians. Elevated PD mortality in post-secondary teachers in the US was reported [6], in white female teachers in particular [12]. There was a suggestion of excess of PD in primary, but not college or secondary, female teachers, based on 17 exposed cases in a population-based case-control study in the US [14]. Conversely, more bias-prone population-based case-control studies in the US [8] and Canada [10] do not support association with work in education and healthcare. Nonetheless, there was a suggestion of an elevated risk for case-control studies for lab technicians in healthcare industry (unlike nursing and other healthcare occupations) after control for referral bias, albeit based on only 15 exposed cases [10], and in female medical/dental technicians (but not in nurses), 9 exposed cases [14]. Resonating with our findings, social work, defined broadly, has been associated with PD in studies of diverse designs [6 8 10], including excess among white female religious workers [12] and evidence that women with PD are more likely to have history of work that requires complex inter-personal interactions [15]. Our observation of tendency towards elevated risk among building and grounds cleaning and maintenance occupations is consistent with exposures to microbes, pesticides, solvents, and cleaning agents, but is not specific. Curiously, there is a suggestion of elevated risk among “janitors/cleaners” in a case-control study from Canada [10]. Tendency towards higher risk with duration of farm living is consistent with current synthesis of evidence [2] and reinforces findings from male-dominated samples. Comparison to literature is hampered by lack of reports of sex-specific analyses and absence of occupation-specific risk estimates in many reports.

In interpreting our findings, we need to consider the selection of women into the WHI-OS and other selection biases. Recruitment into the WHI-OS had no regard for occupation or PD risk, and yet selection on the basis of socio-economic status and expected longevity is a concern with respect to external and internal validity of occupational findings. Of greater concern is whether this cohort represents working conditions of women in modern times, especially for job affected by technological innovation were we see evidence of excess risk, such as medical professions and cleaning. Selection into jobs with low demand on manual dexterity, such as those from retail to counselling or social work due to sub-clinical manifestations of PD is a concern that cannot be addressed by the WHI-OS data; however the same selection mechanism is implausible for top executives.

Outcome misclassification is a worry in epidemiology of PD due to the complexity of the diagnosis. However, we are reassured that self-report proved to be accurate in reliability studies [22-24]. Assessment of the majority of PD cases after interview about work history minimizes odds of differential outcome misclassification with respect to exposure. However, we are still left with the possibility of differential diagnosis by occupation, although (a) our prospective population-based design helps guards against this [10] and (b) we see no consistent excess of risk among medical professionals (who can be presumed to be attuned to early and accurate diagnosis and its recall). Of greater concern is the likely heterogeneity of PD etiologies, whereby different pathologies and causes manifest as a single clinical entity. If this is true, it degrades the power to detect risk due to a single exposure, whose effect size may be modified by the peculiarities of the population (heredity, access to care, etc.) and competing causes.

Our data has limited utility for addressing hypothesis regarding exposures that appear to have been historically uncommon among US women, such as manganese, industrial solvents and organophosphate pesticides, as can be judged on the basis of occupations. However, it is possible to speculate about the role of light-at-night that was implicated in prior work [1], since evidence for effects of this exposure (e.g. in breast cancer epidemiology) rests almost exclusively with women’s work as nurses [29]. In this respect, lack of association with duration of employment as a nurse (consistent with prior evidence [7 10 14]) can be seen as reassuring. Likewise, exposure to anesthetic gasses, such as trichloroethylene [4], may be inferred for some health technologists and technicians for whom there was evidence of elevated risk with longer duration of work, presumably entailing higher historic exposures.

Errors in exposure assessment with duration as proxy for biologically relevant metrics is a concern [30], as is the fact that we did not have full occupational histories. The chance of such errors being differential with respect to PD outcome is guarded against by prospective collection of the majority of the outcome data. However, we cannot rule out that errors in exposure assessment are differential with respect to exposure (e.g. either due to dichotomization of exposure on the basis of occupational code and categorization of estimated duration of work [31 32], or due to chance [33]). Even if departure from non-differential misclassification is negligible, it could have had an effect of both masking true associations and generating spurious ones [34 35]. Nonetheless, we are as confident as one can be in coding of occupations, which was performed using the best existing practices and relied on data of typical quality, and thus can give credence to observed association with occupations, even if any speculation about underlying causal exposure must necessarily remain tenuous.

## Conclusion

Our findings paint a largely reassuring picture of occupational risks for PD among US women, especially for trades largely unaffected by recent technological advances. Nonetheless, more in-depth attention to PD risk in occupations in health technology, social work, teaching, and building & ground cleaning & maintenance may lead to etiologic insights. Our results have no direct policy implication, because we did not assess specific exposure and make no causal claims. However, using strong design, reliable methods, and a large sample, we succeeded in producing finding that can help target the search for occupational causes of PD among women.

## Authors’ contributions

IB and HC participated in conception and design of the work; IB, AZL, RBW and HC contributed to the acquisition of the data; all authors participated in analysis, interpretation of data for the work, as well as drafting the work. All authors agree to be accountable for all aspects of the work in ensuring that questions related to the accuracy or integrity of any part of the work are appropriately investigated and resolved.

Authors’ contributions
IB and HC participated in conception and design of the work; IB, AZL, RBW and HC contributed to the acquisition of the data; all authors participated in analysis, interpretation of data for the work, as well as drafting the work. All authors agree to be accountable for all aspects of the work in ensuring that questions related to the accuracy or integrity of any part of the work are appropriately investigated and resolved.

## Acknowledgements

The authors are thankful to Dr. Loni Philp Tabb for helping shape approach to statistical analysis. Furthermore, for the current manuscript, Ms. Ruoxuan Qi assisted with SAS programming and Dr. Angla Wang contributed to review of cases identified by use of medication.

